# A full-length infectious cDNA clone of a dsRNA totivirus-like virus

**DOI:** 10.1101/2021.11.08.467288

**Authors:** Han Wang, Kenta Okamoto

## Abstract

Totivirus-like viruses are icosahedral non-enveloped double-stranded (ds)RNA viruses belonging to a group recently discovered and provisionally assigned in the *Totiviridae* family. Unlike fungal and protozoan *Totiviridae* viruses, these totivirus-like viruses infect a diverse spectrum of metazoan hosts and currently have enormous impacts on fisheries and agriculture. We developed the first totivirus-like virus Omono River virus (OmRV) infectious full-length DNA clone and produce the infectious particles using an RNA-transcript-based method. Unlike the parent wild-type particles from nature, the reverse-genetically-generated OmRV particles had an indistinguishable cytopathic effect, infectivity, and morphology. The established system is one of the few systems that have been reported for generating a non-segmented dsRNA virus DNA clone.

## Introduction

Icosahedral non-enveloped double-stranded (ds)RNA viruses are currently infecting a wide range of hosts, from prokaryotes to eukaryotes. Notably, these viruses have enormous impacts on fisheries, animal husbandries, food manufacturing and even on human health worldwide [1– 6].

*Totiviridae*, one of the taxa in icosahedral non-enveloped dsRNA viruses, encompasses five approved genera: *Giardiavirus, Leishmaniavirus, Totivirus, Trichomonasvirus* and *Victorivirus* [7–9]. Their virions are approximately 40 nm in diameter and are composed of single 4.6-7.0 kbp in-size dsRNA genomes encoding one major capsid protein (CP) and an RNA-dependent RNA polymerase (RdRp) [10, 11]. Few of them encode additional functional proteins [12]. The viruses in the *Totiviridae* family were previously reported to be infectious on yeast, smut, filamentous fungi and parasitic protozoa [13]. A new group of totivirus-like viruses that infects metazoan hosts has been discovered of late, but the viruses have been provisionally placed in the *Totiviridae* family and have not been definitively classified. The totivirus-like viruses have so far been isolated from arthropods such as shrimps [1]and insects (e.g. mosquitoes, flies, ants, mosses) [10, 14-16] and from vertebrates such as fishes [11, 12], including the Omono River virus (OmRV) in this study [15]. Another group of dsRNA viruses, *Tianjin totivirus* (ToV-TJ), was extracted from bat feces [17], and from tea oil trees in China [18]. Some of these viruses greatly and ceaselessly affect fishery cultivations around the world [1, 12]. Their ability to infect a variety of metazoan hosts has drawn much attention to them. In the previous studies, it was hypothesised that the capsid of the totivirus-like viruses should have acquired several unique functions in cell entry and gene replication, which are tied into their survival strategies in multicellular hosts [7, 8, 19]. Hence, the first infectious clone of the totivirus-like viruses will help in understanding their molecular mechanisms with regards to their unique functions and pathogenicity.

For RNA viruses, the commonly used reverse genetic platforms include the syntheses of full-length genomic RNA transcripts (the RNA-transcript-based method), full-length cDNA/RdRp products or bacterial artificial chromosome (BAC)/RdRp, followed by the transfection or electroporation into host cells. These approaches have been applied in a variety of positive-sense single-stranded ((+)ss)RNA viruses [20–27]. For segmented dsRNA viruses, infectious clones of *Reoviridae* viruses obtained using the RNA-transcript- and plasmid-based methods [14, 28, 29–36, 37-39]. For non-segmented dsRNA viruses, however, few successful cases have been obtained using the RNA-transcript-based method [40, 41]. We first generated an infectious totivirus-like virus from a constructed full-length 7.6 kbp OmRV infectious clone using an RNA-transcript-based method, which will be great additions for studying their molecular mechanisms.

## Materials and Methods

### Infectious cloning of Omono River virus and seed virus preparation

OmRV-fragment1/pMK and OmRV-fragment2/pMA plasmids designed on the basis of the full genome sequence of the OmRV AK4 strain (GenBank accession number: AB555544) were ordered from GeneArt Custom Gene Synthesis Service (Thermo Fisher Scientific). OmRV-fragment1/pMK (kanamycin resistant), OmRV-fragment2/pMA (ampicillin resistant) and pACYC177 low-copy-number plasmids were transformed into 10G chemically competent cells (Lucigen, E. cloni), respectively. The OmRV-fragment1/pMK contains an SP6 promoter sequence just before the OmRV gene. The transformed *Escherichia coli* cells were propagated at 25°C, 150 rpm for 48 h. Sufficient amounts of these three plasmids (> 1 µg/µL) for cloning were purified from these transformed *E. coli* cells using PureLink HiPure Plasmid Maxiprep Kit (Thermo Fisher Scientific). Then OmRV-fragment1/pMK was digested with DraIII and XhoI at 37°C overnight. The digested product was then electrophoresed on 1% (w/v) agarose gel and purified from the gel slice using PureLink Quick Gel Extraction Kit (Thermo Fisher Scientific). The purified product was subcloned into pACYC177 plasmid at DraIII-XhoI sites. The reconstructed product OmRV-fragment1/pACYC177 was then transformed into 10G chemically competent cells (Lucigen, E. cloni), propagated for plasmid purification with the aforementioned method. Fragment 2, which was digested from OmRV-fragment2/pMA by XhoI and Cfr9I at 37°C overnight was subcloned into OmRV-fragment1/pACYC177 at XhoI-Cfr9I sites. Consequently, the OmRV-full-length/pACYC177 was cloned and used for synthesising viral RNAs. The finally constructed OmRV-full-length/pACYC177 infectious clone (OmRV/IC) is shown in Figure 1. The reconstructed OmRV/IC plasmid was then transformed into electrocompetent *E. coli* (Lucigen) through electroporation (2.5 kV, 5 ms) and the cells were spread on a Luria-Bertani (LB)-agar plate (Miller, Sigma-Aldrich) supplemented with 100 µg/mL ampicillin. The plate was placed and incubated at room temperature for 2 days, until the colonies formed. The transformants were then propagated in an LB liquid medium with 100 µg/mL ampicillin at 25°C, 150 rpm for 48 h. The purified product was linearised with 1 h digestion of XhoI, and then extracted from electrophoresed 1% (w/v) agarose gel (SYBR™ Safe DNA Gel stain, Thermo Fisher Scientific), followed by DNA sequencing. No mutation was detected in the OmRV/IC plasmid.

**Figure 1.**
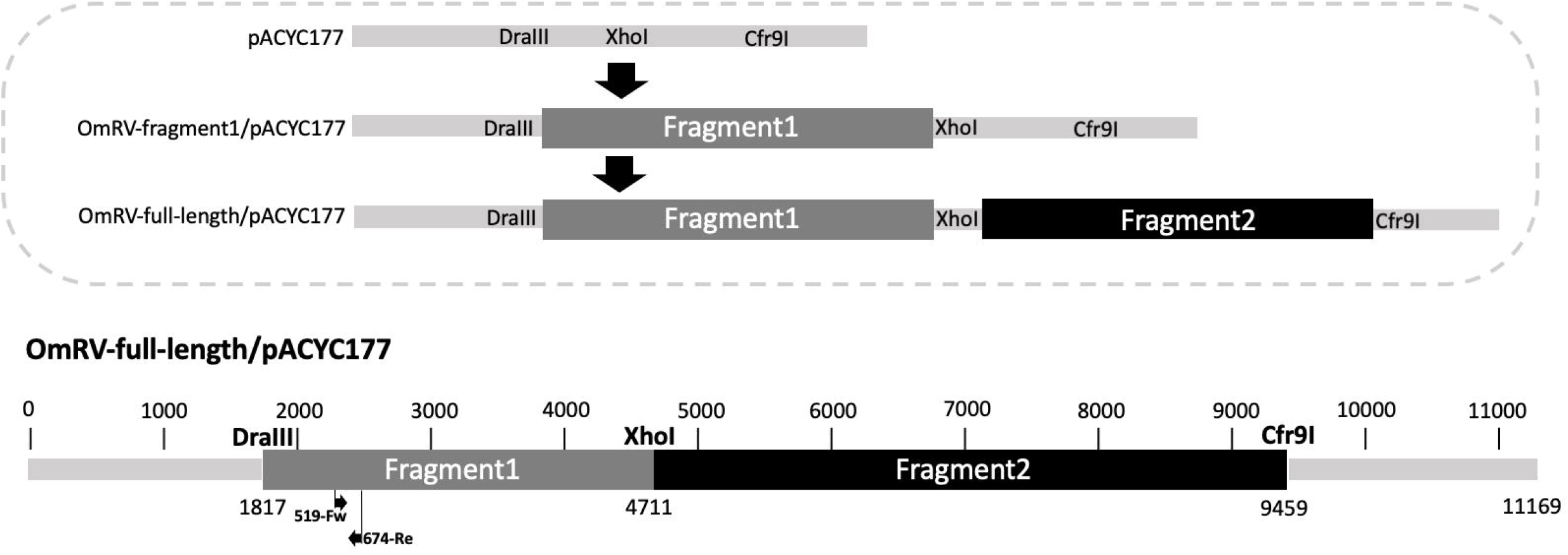
Schematic description of designing and cloning OmRV cDNA infectious clone using a low-copy-number plasmid pACYC177. The upper panel shows the process of cloning from the beginning to the end. The bottom is the full construct of the designed OmRV-full-length/pACYC177. The primer set 519-Fw and 674-Re for viral RNA detection is also included and pointed out with black arrows. SP6 promoter gene sequence is located just before the OmRV gene of the fragment 1 to initiate the RNA synthesis.

For in vitro transcription, the OmRV/IC plasmid (10 µg) was linearised with Cfr9I enzyme and transcribed to RNA with mMESSAGE mMACHINE High-Yield Capped RNA Transcription SP6 Kit (Thermo Fisher Scientific) following the manufacturer’s instructions. OmRV/IC RNA (2.5 µg) was introduced into 5 × 10^6^ C6/36 mosquito cells per well of a 6-well plate via Lipofectamine 2000 reagent (Thermo Fisher Scientific), following the product’s instructions. The same amount of Lipofectamine 2000 reagent-treated C6/36 cells was also used as a negative control. Both were prepared in triplicate in three wells. 200 µL of the infected culture fluid (ICF) was then collected on 0, 1, 3, 5, and 8 days post-infection (dpi), respectively. The viral RNA from the ICF samples were extracted using PureLink Viral RNA/DNA Mini Kit (Thermo Fisher Scientific) following the corresponding instructions.

### Virus titration using quantitative reverse transcription polymerase chain reaction

A primer set 519-Fw (5’-TGTGTATAAGGTTGGGTCGGAAG-3’) and 674-Re (5’-GACAACAAACACATAGGACAGAA-3’) specifically for a part of the fragment 1 region of OmRV was used in quantitative reverse transcription polymerase chain reaction (RT-qPCR) for confirming the progeny recombinant OmRV particle propagation (Figure 1). The viral RNA level was determined according to the standard samples. The standard DNA construct [Std-OmRV (508-684)/pMA] that is composed of an SP6 promoter, OmRV-AK4 (508-684) and an XhoI restriction site in a pMA plasmid, was designed and ordered as for the RT-qPCR standard. The 5 µg of Std-OmRV (508-684)/pMA plasmid was linearised with an overnight digestion of XhoI at 37°C and was then transcribed to RNA using mMESSAGE mMACHINE High-Yield Capped RNA Transcription SP6 Kit (Invitrogen) for the following RT-qPCR. All the samples containing viral RNA were extracted from the same volume of ICF. Power SYBR Green PCR Master Mix (2X), MultiScribe Reverse Transcriptase (50 U/mL) and RNase Inhibitor (20 U/mL) (Thermo Fisher Scientific) were also used as recommended in the manufacturer’s instructions. The RT-qPCR conditions were as follows: 48.0°C for 30 min, 95.0°C for 10 min, followed by 40 cycles of 95.0°C for 15 s and 60.0°C for 1 min. The total reaction volume in each well was 10 µL. The C_T_ values and other parameters were analysed, generated and exported from the system software (QuantStudio™ 6 Flex System). The data were then processed and the figures were generated from the Graphpad Prism 8 software according to the standard curve.

### Cell culture

C6/36 Aedes mosquito cells were provided from Morita’s group at NEKKEN, Nagasaki University, and were cultured in a minimum essential medium (Eagle, Sigma-Aldrich) supplemented with non-essential amino acid (NEAA, Gibco), 10% (v/v) fetal bovine serum (FBS, Biowest) and 100 U/mL penicillin/100 µg/mL streptomycin (Gibco) at 28°C in a 5% CO_2_ atmosphere.

### Plaque assay

After two passages of OmRV/IC ICF in C6/36 mosquito cells, an aliquot of the OmRV/IC containing ICF was collected. Plaque assay was then performed for testing the infectivity of the generated OmRV/IC particles. The OmRV-AK4 parent virus (OmRV/Ori) from nature [15] was also used as a positive control. The 1 mL of each of the serially diluted OmRV/IC and OmRV/Ori samples was prepared and added in 90-100% confluent monolayer C6/36 mosquito cells in each well of the 6-well plates. Each serial dilution was tested in triplicates. The plates were incubated at 28°C, 5% CO_2_ with gentle rocking for even coverage and to prevent them from drying out. After 1 h incubation, the fluid containing virus particles was gently removed, and 2 mL of the freshly prepared sterile immobilising MEM medium supplemented with NEAA, 2% (v/v) FBS, 0.4% (w/v) agarose, and 100 U/mL penicillin/100 µg/mL streptomycin was overlaid on the inoculums in each well at room temperature. The plates with overlays were then incubated at 28°C, 5% CO_2_, for 3-5 days, until clear plaques appeared. The C6/36 mosquito cells were directly fixed in each well with the addition of 1 mL of 4% (v/v) paraformaldehyde directly in each well and were kept at room temperature for 1 h. Then the paraformaldehyde solution and immobilising overlays were removed, and the wells were washed with ddH_2_O twice. The fixed C6/36 mosquito cells were then stained with 1% (w/v) crystal violet (Sigma-Aldrich) dissolved in 20% (v/v) ethanol.

### Virus particles propagation, isolation, purification and electron microscopy imaging

To obtain a higher yield of the OmRV/IC particles, the seed OmRV/IC particles were propagated in five 175 cm^2^ flasks. The cells were incubated until they were detached due to the strong cytopathic effect (CPE). The OmRV-IC/wt particles were isolated and purified through a previously reported method [7]. The viral proteins were detected by SDS-PAGE. The purified OmRV/IC particles were loaded onto a glow-charged carbon-coated grid (TED PELLA, 01753-F) and were negatively stained using gadolinium salts. The grid was observed using a transmission electron microscope (FEI Technai G2) under an 80 kV acceleration voltage.

## Results and Discussion

The non-enveloped icosahedral dsRNA viruses synthesise their positive-sense single-stranded RNAs ((+)ssRNAs) genome intrapartically. The (+)ssRNAs, templates for synthesising viral capsid protein (CP) and RNA-dependent RNA polymerase (RdRp), assemble into a replication intermediate that is composed by CP and RdRp. In turn, the dsRNA is synthesised by the (+)ssRNA template and RdRp in the assembled capsid [42]. The dsRNA totivirus-like viruses are expected to employ a similar mechanism for intraparticle genome synthesis [8]; thus, reverse-genetically-synthesised OmRV/IC (+)ssRNA was introduced into the C6/36 mosquito cells for infectious-particle generation.

The OmRV RNA copy number kept increasing from day 1 to day 8 after the OmRV/IC RNA was introduced into C6/36 mosquito cells (Figure 2A), demonstrating the virus propagation from the synthesised viral RNA. Although the propagated viruses showed no or weak CPE in the RNA-transfected C6/36 cells, the virus titres went up remarkably, displaying significant CPE, after several passages of ICF in the C6/36 cells (5.2 × 10^9^ pfu/mL). The OmRV/IC-infected C6/36 cells showed large swelled structures, were detached, and were finally lysed (Figure 2B). The plaques of the infectious OmRV/IC and OmRV/Ori particles were compared (Figure 2C). No obvious differences were seen in the sizes of the plaques (2 mm on average), which indicates that the virus multiplication rates of OmRV/IC and OmRV/Ori are similar. The purified and concentrated OmRV/IC particles were characterised using SDS-PAGE. The results showed several bands, including those of major CP and of other minor proteins (Figure 3A). The transmission electron microscopy images also showed that correctly sized (approximately 40 nm in diameter) OmRV/IC particles were formed (Figure 3B). The OmRV/IC particles were also morphologically indistinguishable from the OmRV/Ori ones.

**Figure 2.**
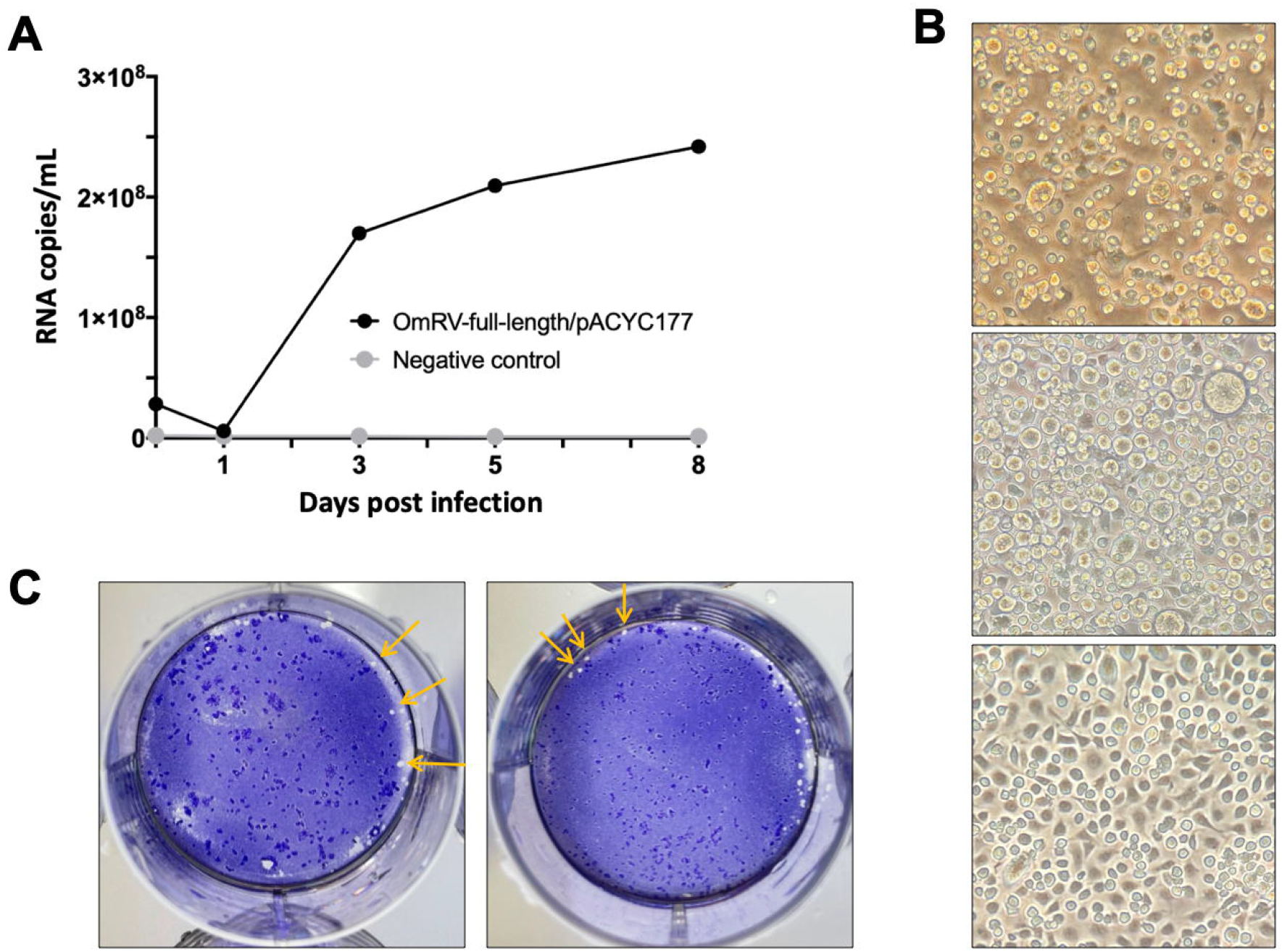
The virus propagation from the OmRV/IC cDNA clone in C6/36 mosquito cells. A) Averaged qPCR titration growth curves of propagated virus particles 0, 1, 3, 5 and 8 days after the introduction of OmRV/IC RNA or its negative control. The number of the RNA copies was calculated from the RT-qPCR results of the standard. The data was shown as an average of the triplicate results. B) The CPE comparisons between the OmRV/Ori (up), OmRV/IC (middle) particles infected C6/36 cells (6 dpi), and healthy C6/36 cells (bottom). C) The plaque assay results of OmRV/IC (left) and OmRV/Ori particles (right) (2 dpi) The plaques are pointed out with yellow arrows.

**Figure 3.**
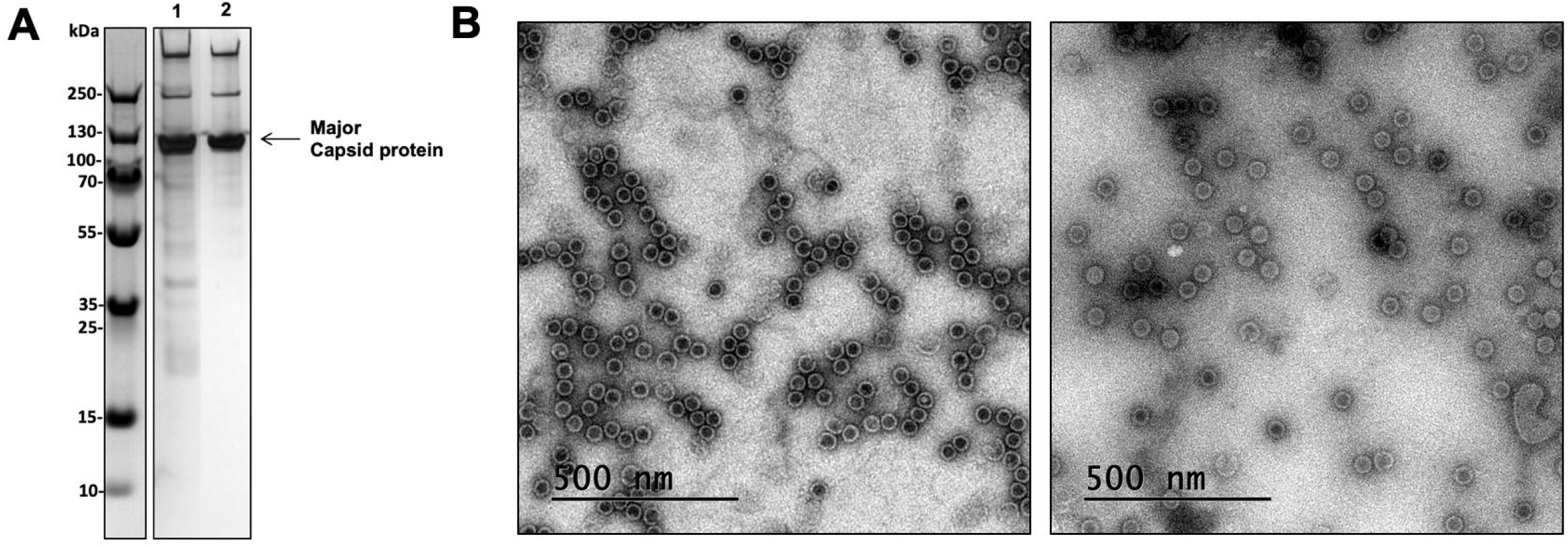
Morphological comparison in OmRV/Ori and OmRV/IC particles. A) The SDS-PAGE result of the purified OmRV/Ori (Lane 1) and OmRV/IC (Lane 2) particles. The black arrow indicates major capsid protein of the OmRV particles. B) The transmission electron microscope images of the purified OmRV/Ori (left) and OmRV/IC (right) particles.

The OmRV/IC is a useful tool for further studies on totivirus-like viruses, including their mutagenesis, biophysical and molecular function and pathogenicity. No significant difference in virus infectivity and morphology was seen between the OmRV/IC and OmRV/Ori particles, which will allow us to produce large amounts of functional OmRV particles and their mutants for extensively clarifying the hypothesised structure-functions of totivirus-like viruses as previously reported [8], [19].

This paper reports one of the few successful cases of generating an infectious DNA clone of the non-segmented dsRNA viruses, and the first case for totivirus-like viruses. Therefore, the described RNA-transcript-based method will also provide ideas for the further design of a DNA clone of other dsRNA viruses that pose threats to agriculture, fishery, and human health.

## Additional information

## Acknowledgements

This work was supported by the following agencies: The Swedish Research Council (VR: Vatenskapsrådet to K.O., grant number: 2018-03387), FORMAS research grant from the Swedish Research Council for Environment, Agricultural Sciences and Spatial Planning (to K.O., grant number: 2018-00421), and The Royal Swedish Academy of Sciences, (KVA: Kungl. Vetenskaps-akademien, Biosciences 2018 to K.O., grant number: BS2018-0053).

Original OmRV AK4 strain was kindly provided by H. Isawa., K. Sawabe. at the Department of Medical Entomology, National Institute of Infectious Diseases (NIID), Japan.

## Author Contributions Statement

H.W., and K.O. designed the experiments, prepared samples, collected and analysed data, and wrote the manuscript.

